# Optoretinographic measurements of cone photoreceptor function in disease-affected retinas

**DOI:** 10.64898/2026.11.16.767206

**Authors:** Reddikumar Maddipatla, Elain Guo, Maciej M. Bartuzel, Yao Cai, Anqi Zhang, Kareem Moussa, Glenn C. Yiu, Robert J. Zawadzki, Ravi S. Jonnal

## Abstract

Optoretinography (ORG) provides a noninvasive approach for measuring stimulus-evoked photoreceptor responses in the living human retina, but its application to retinal disease using clinically translatable imaging systems remains limited. Here, we applied velocity-based ORG using a clinical-grade swept-source optical coherence tomography (OCT) system to assess photoreceptor function in disease-affected human retinas and healthy controls. Following pupil dilation and dark adaptation, repeated B-scans were acquired at 400 Hz during visible-light stimulation using a phase-sensitive 100-kHz, 1060-nm swept-source OCT system. Stimulus-evoked optical path length changes between the inner segment/outer segment junction (IS/OS) and cone outer segment tips (COST) were used to quantify cone outersegment responses. Healthy retinas exhibited rapid and reproducible responses characterized by an initial outer-segment contraction followed by elongation, whereas disease-affected retinas exhibited quantitatively reduced or altered response amplitudes and temporal dynamics. ORG also detected functional improvement following treatment in a vitamin-A-deficient retina. These findings demonstrate the ability of velocity-based ORG to quantitatively detect photoreceptor dysfunction and functional change using a clinically compatible OCT platform, supporting its potential as a noninvasive functional biomarker for retinal disease and therapeutic response.

## 2 Introduction

In blinding retinal diseases such as age-related macular degeneration (AMD) and retinitis pigmentosa (RP), dysfunction of photoreceptors is thought to precede their outright loss. Currently, in ophthalmic clinics, these diseases are detected and assessed by observing structural anomalies of the retinal layers using optical coherence tomography (OCT), normally in combination with subjective tests such as visual acuity (VA) and visual fields (VF). An ideal complement to these techniques would be an objective functional test based on imaging of the photoreceptors, which would reveal functional and structural changes simultaneously. Objective assessment of the functional response could greatly help ophthalmologists detect retinal diseases at an early stage and manage them effectively. It could also catalyze trials of therapeutics by supplying them with earlier and more sensitive endpoints.

OCT is a non-invasive, high-resolution, volumetric imaging technique commonly used to assess retinal layer abnormalities. In OCT the amplitude and phase of the light reflected by the sample are recorded, and usually the amplitude of the signal is used to generate high-resolution structural images of the sample, while the phase is ignored. By utilizing the phase of the OCT signal, nanometer-scale changes in the sample can be quantified,^1^ which offers the ability to measure subtle structural changes in the photoreceptors due to external light stimuli.

Early efforts to observe these stimulus-evoked responses in human cone photoreceptors used adaptive optics (AO), enabling resolution and tracking of single cells, in conjunction with common path interferometry.^2^ Since then, interest in this area has grown, and these responses have been successfully measured in single cones^3–7^ and rods^8^ using advanced OCT systems with cellular resolution achieved with adaptive optics (AO) or digital aberration correction (DAC). Full-field swept-source OCT has been used to measure responses to diverse stimuli with AO^9^ and to chirped flicker stimuli with DAC.^10, 11^ These approaches measure responses from single cells, most commonly by monitoring the phase difference between outer segment (OS) tips of cones (COST) or rods (ROST) and the inner-outer segment junction (IS/OS), and this approach has been termed optoretinography (ORG). Two key features of the photoreceptor response to light are (1) an initial OS contraction lasting 10 ms to 20 ms and (2) an ensuing OS contraction lasting ≥100 ms, which have now been observed in more than ten labs around the world.

AO enables cellular ORG measurements because it permits the tracking of single cells over time. In this way, the “purest” responses are recorded, i.e., those without artifacts due to extracellular factors. It also permits complementary analysis of cell properties and structure both in health and disease.^12, 13^ The AO-ORG systems described thus far have been extremely fast, capitalizing on full-field OCT,^3^ megahertz+ swept sources,^4, 8^ spectrometer multiplexing,^5^ or line-illumination with 2D detection.^6^ By being fast they were able to achieve both cellular resolution and the necessary temporal resolution to detect the key features of the photoreceptor ORG response. However, measuring ORG signals with an advanced AO system poses some challenges for clinical translation. Research-grade AO-OCT systems are costly and typically require multiple expert operators. The volume of processed data can be in the terabytes for a single experiment, requiring costly storage and backup systems, as well as protracted data processing times.

While most recent work in this area has utilized AO and the phase of the OCT signal, some investigators have shown that the amplitude (and speckle) of the OCT signal can be used to measure light-evoked changes in the retinas of animal models^14–16^ and humans.^17, 18^ Additional detail on these and other ORG methods can be found in recent reviews.^19, 20^

A challenge posed by clinical-grade OCT instrumentation is its relatively low speed. To measure the fastest ORG dynamics, volumetric frame rates should be over 100 Hz, which are afforded only by the uncommon high-speed OCT implementations listed above. Typical research-grade OCT implementations and clinical OCT systems cannot achieve volume rates this high, but can generally achieve sufficient B-scan rates. However, implementing the ORG with serial B-scanning requires a paradigm shift from the AO-ORG, since in awake humans the retina cannot be expected to be stationary for the ∼50 ms required to measure the initial contraction and ensuing elongation of the OS. Thus, over the course of measurement, different parts of the retina are imaged. In the AO-ORG paradigm the phase of light reflected by structures is monitored over time, as those structures are tracked in three dimensions. In the absence of tracking, the structures in view vary over time. If this paradigm were applied to serial B-scans, microscopic axial variations in the structures being imaged, as they moved in and out of the scanning beam, would result in spurious phase changes mistaken for ORG signal.

We recently published results from a novel ORG paradigm designed for an OCT platform very similar to those used in clinics.^21^ The method works by computing the relative velocities of the IS/OS and COST over a brief (5 ms to 10 ms) window and rolling that window over the series of B-scans acquired over much longer durations. The stimulus flash must be larger than the B-scan’s scan line to account for small fixational eye movements during those longer durations. The key assumptions of this paradigm are 1) that photoreceptors that are near one another exhibit correlated ORG responses; 2) there is a finite interval over which eye movements are negligible compared to the spot size of the OCT; and 3) changes in the *rate* of OS length change are commensurable with changes in the OS length. In the present work, we explore this method’s ability to measure deficits in photoreceptor function in patients with vision loss. We selected patients with a variety of retinal disorders in order to demonstrate broadly the method’s sensitivity to disease-related dysfunction and motivate further investigation of specific diseases.

## 3 Methods

### 3.1 Imaging system

ORG imaging was performed using a phase-stable swept-source OCT system described previously.^21, 22^ Breiefly, the system consisted of a 100 kHz swept-source laser (Axsun; *λ* = 1060 nm; Δ*λ* = 100 nm), two galvanometric scanners (Cambridge), a transmissive reference arm, a balanced photoreceiver (Thorlabs), and a digitizer (Alazar).

The power of the imaging source, measured at the cornea, was 1.8 mW.^21–23^ A fibercoupled 555 nm LED stimulus was synchronized with OCT acquisition to evoke photoreceptor responses. Repeated B-scans were acquired at 400 Hz, and optical path length changes between IS/OS to COST of cone photoreceptor were computed using previously reported velocity based phase-sensitive methods.^21, 22^

Additional details regarding system configuration, stimulus delivery, and signal processing are provided in previous publications.^21–24^

### 3.2 Human subjects

Sixteen subjects were imaged in this study, including six healthy controls and ten patients with retinal pathology. The patient cohort consisted of two subjects with macular telangiectasia type 2 (MacTel), one with diabetes without retinopathy, one with diabetic retinopathy, two with unexplained bilateral vision loss, two with retinitis pigmentosa (RP), one with Stargardt maculopathy, and one with vitamin A deficiency.

Healthy controls (age range, 21–50 years) had no known systemic or ocular disease and included one female and five males. Patients ranged in age from 49 to 75 years.

All subjects provided written informed consent prior to participation. The study was approved by the Institutional Review Board (IRB) of the University of California, Davis, and adhered to the tenets of the Declaration of Helsinki. Subjects were referred to by anonymized, IRB-assigned identification numbers that are unique across multiple ongoing studies in our laboratory; thus their numbering is not consecutive within this study.

Subject demographics and diagnoses are summarized in Table 1.

**Table 1:** Subject demographics and diagnosis.

| Subject ID | Age | Sex | Diagnosis |
|---|---|---|---|
| NR | 21–50 | 1F/5M | Healthy controls |
| 23 | 58 | F | Stargardt maculopathy |
| 27 | 75 | M | Unexplained vision loss |
| 28 | 61 | M | Unexplained vision loss |
| 30 | 49 | F | Vitamin A deficiency (VAD) |
| 33 | 48 | F | Retinitis pigmentosa |
| 34 | 50 | F | Retinitis pigmentosa |
| 52 | 62 | M | MacTel Type 2 |
| 69 | 60 | F | MacTel Type 2, diabetes w/o retinopathy |
| 60 | 58 | F | Diabetic retinopathy (T2D) |
| 61 | 72 | F | Diabetes without retinopathy |

Prior to imaging, anterior chamber angles and intraocular pressure were measured to ensure a low risk of acute angle-closure glaucoma following dilation. The eyes were dilated using topical drops of tropicamide 1% and phenylephrine 2.5%.

### 3.3 Imaging protocol

Subjects were dark-adapted for five minutes prior to imaging. At each retinal location, a series of 80 B-scans was acquired over 200 ms, with the stimulus flash presented 100 ms after the start of acquisition. To facilitate comparisons across subjects, ORG measurements were acquired at five retinal eccentricities: 2*^◦^*, 4*^◦^*, 6*^◦^*, 8*^◦^*, and 10*^◦^* relative to the foveal center. At each eccentricity, five measurements were obtained at different locations along an iso-eccentric arc centered on the fovea, with the polar angle varied while maintaining a constant radial distance. The locations were spaced to avoid overlap between sucessive stimulus exposures. Thus, a total of 25 ORG measurements were acquired following a single dark-adaptation period. Including dark adaptation, the imaging session required less than ten minutes, excluding subject alignment and OCT system optimization. Further details of the imaging protocol have been reported previously.^21–24^

### 3.4 Swept-source OCT signal processing

OCT B-scans were processed using standard swept-source OCT signal processing procedures, similar to previously reported methods.^22, 25^ Raw spectral interferograms acquired during each wavelength sweep were first cropped to remove noisy regions at the beginning and end of the sweep. The DC component of the signal was estimated by averaging spectra from fast scans and subsequently subtracted from each acquired interferogram. Nonlinear sampling in wavenumber and group velocity dispersion mismatch between the sample and reference arms introduced phase chirp in the interferograms. These effects were corrected numerically by applying a polynomial phase correction. The coefficients of the dechirping polynomial were selected through an optimization algorithm. Following these corrections, discrete Fourier transformation (DFT) was applied to convert the corrected interferograms into complexvalued B-scans. The resulting B-scans were stored as complex-valued arrays for subsequent phase-sensitive analysis.

### 3.5 Calculation of rate of outer segment length change (*v_OS_*)

The rate of outer segment length change (*v_OS_*) was computed using a velocity-based optoretinography method described previously.^22, 25^ Complex B-scans were registered and grouped into overlapping blocks of three consecutive frames acquired in 7.5 ms. Withing each block, bulk axial motion was corrected numerically,^26^ followed by semi-automated segmentation of the IS/OS and COST layers. Detailed definitions of the segmented retinal layers are provided in a previous publication.^23^ Phase differences between segmented layers were computed at each lateral location and time point, expressed as Δ*θ_n_*(*x, t*). The least-squared error linear fitting of the unwrapped phase with respect to time yielded phase velocity estimates, *v_OS,n_*(*x*). These velocities then averaged across the lateral dimension over the stimulus region to obtain the rate of outer segment length change, *v_OS,n_* for each block.

A schematic of the processing pipeline is shown in Fig.1. Additional implementation details are provided in a previous publication.^22, 25^

**Figure 1:**
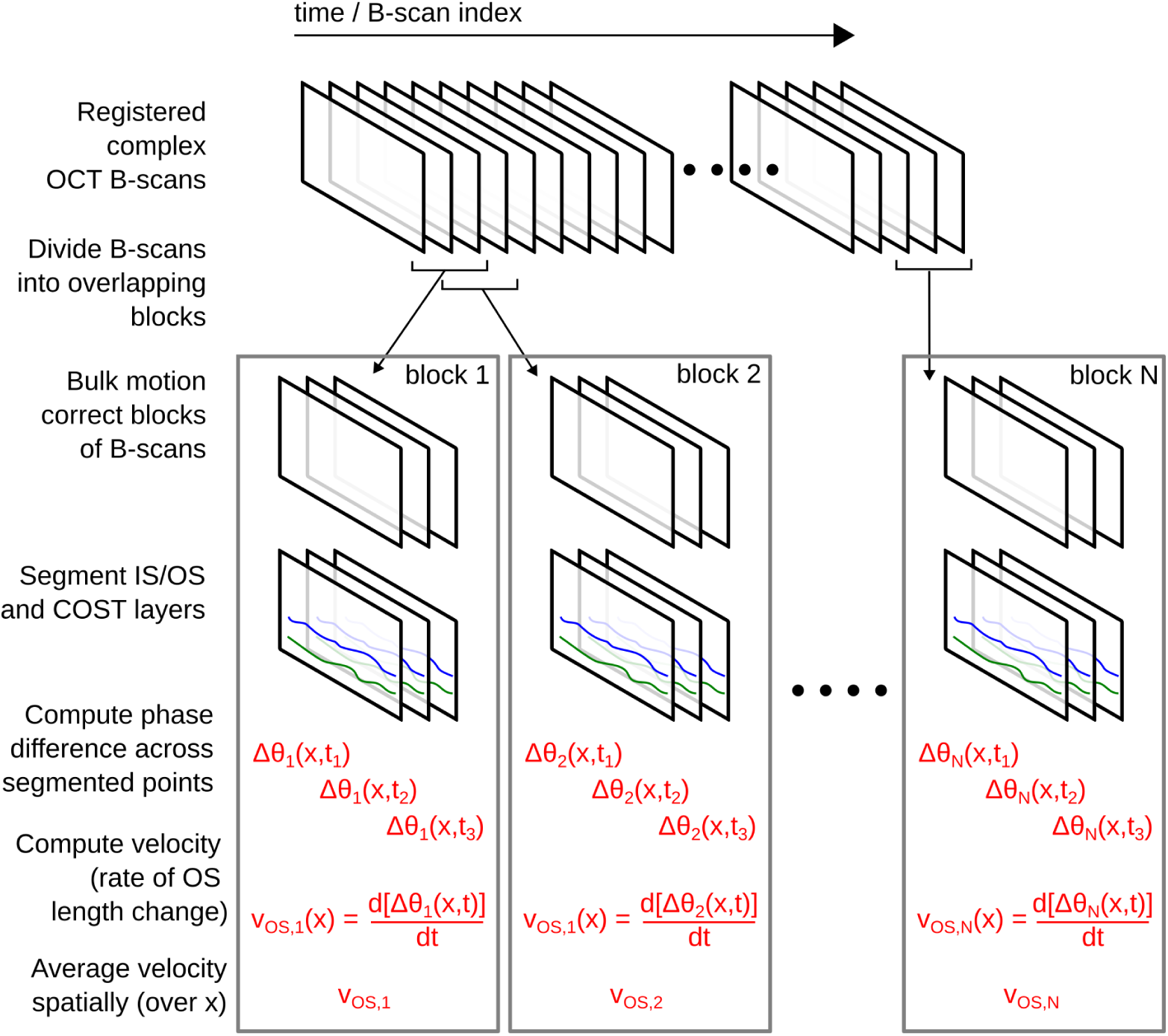
Velocity-based ORG data processing steps. Complex B-scans were aligned and divided into overlapping blocks of three. B-scans in each block were bulk-motion corrected, and the IS/OS and COST layers were segmented. In each B-scan, at each pair of segmented IS/OS and COST coordinates, the phase difference was calculated: Δ*θ_n_*(*x, t*), where *n*, *t*, and *x* represent the block index, time, and location along the fast-scanning dimension. Next the slope of the phase was determined at each *x* coordinate, resulting in *v_OS,n_*(*x*). These phase velocities were then averaged spatially to produce each block’s rate of OS length change, *v_OS,n_*.

### 3.6 Quantification of ORG responses

ORG responses were quantified by first averaging measurements across the five locations at each eccentricity to obtain a mean response for each subject. Two previously described parameters^22, 25^ were used to characterize the response amplitude:

1. *v_min_* was defined as the most negative value observed after the flash.
2. *v*_20_*_−_*_40_ was defined as the mean velocity over the 20 ms to 40 ms interval following stimulus onset.

These parameters were computed at each eccentricity for all subjects. To facilitate comparison across subjects, patient values were normalized by the corresponding values obtained from control subjects.

## 4 Results

### 4.1 Representative OCT images and ORG responses in healthy and disease-affected retinas

The OCT B-scans from all subjects and retinal eccentricities exhibited clearly identifiable IS/OS and COST bands, allowing reliable segmentation and subsequent ORG analysis using the semi-automated procedure described above. Representative OCT B-scans and corresponding ORG responses from a healthy control subject (#16) and a patient with vitamin A deficiency (#30) are shown in Fig. 2. The highlighted regions in the B-scans indicate the locations used for ORG analysis. Measurements from the vitamin A-deficient subject were acquired before and after vitamin A supplementation treatment.

**Figure 2:**
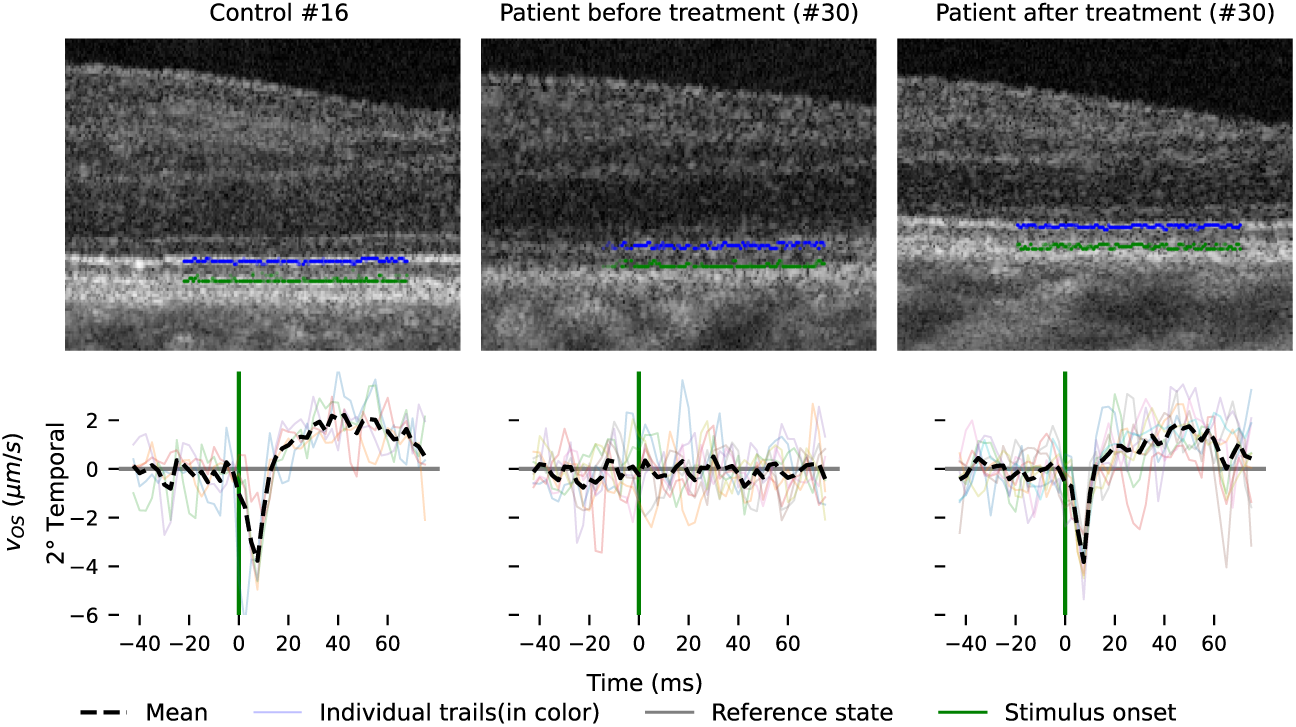
Representative OCT B-scans and corresponding ORG responses from a healthy control subject and a patient with vitamin A deficiency. Top row: representative OCT B-scans from (a) a healthy control retina (#16), (b) vitamin A deficiency (#30) before treatment, and (c) after vitamin A supplementation treatment. The ORG responses were computed from the points highlighted in the IS/OS and COST bands. Bottom row: corresponding ORG responses *v_OS_* from the same retinal location. The healthy control exhibited a characteristic biphasic response consisting of rapid contraction followed by elongation. Prior to treatment, the vitamin A deficient subject exhibited attenuated ORG responses, particularly reduced post-stimulus elongation. Following treatment, recovery of the ORG response was observed.

In the healthy control subject, the ORG response demonstrated the characteristic biphasic profile observed in all normal subjects, consisting of an initial rapid contraction upon the stimulus onset followed by a slower elongation. Prior to treatment, the vitamin A deficient subject exhibited a substantially attenuated ORG responses compared with the healthy control, despite relatively preserved outer retinal structure seen in the OCT images. Both the initial contraction velocity (*v_min_*) and the elongation velocity between 20 ms to 40 ms (*v*_20−40_) are nearly absent.

Following vitamin A supplementation, the ORG response exhibited substantial recovery toward normal response characteristics, including increased elongation amplitude and improved temporal dynamics. These results demonstrate that ORG can detect functional abnormalities and recovery in disease-affected retinas, even when the structure of the retinal layers looks normal in OCT cross-sectional images.

### 4.2 ORG responses in healthy controls

ORG responses from healthy control subjects were measured at retinal eccentricities ranging from 2*^◦^* to 10*^◦^* (Fig. 3). Response amplitudes generally decreased with increasing eccentricity, although inter-subject variability was observed. Despite these variations, healthy subjects demonstrated reproducible responses across retinal locations and eccentricities.

**Figure 3:**
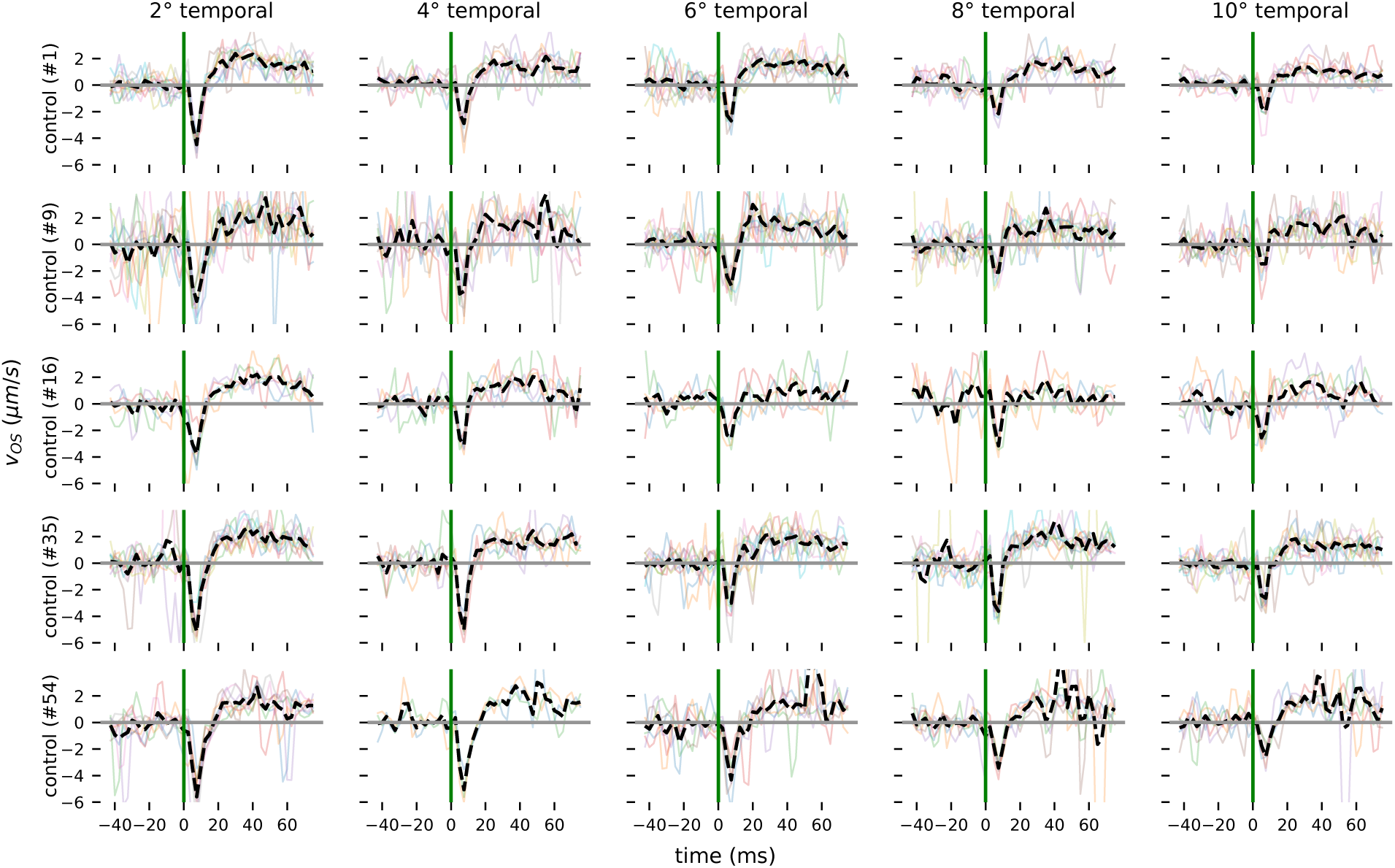
Representative ORG responses from healthy control subjects. ORG responses from the healthy control subjects at retinal eccentricities ranging from 2*^◦^* to 10*^◦^*. Colored traces represent individual measurements acquired from iso-eccentric retinal locations, while dashed black traces indicate the averaged response at each eccentricity. The vertical green lines indicate stimulus onset, and the horizontal gray line indicates zero velocity. Measurements exhibiting excessive motion artifacts were excluded from averaging and display. Healthy subjects consistently exhibited a characteristic stimulus-evoked ORG response across retinal eccentricities.

### 4.3 ORG responses in vitamin A deficiency

The vitamin-A-deficient subject (#30) exhibited markedly attenuated ORG responses compared with healthy control subjects (Fig. 4). Prior to treatment, both the early contraction and subsequent elongation phases were substantially reduced across retinal eccentricities. Following intramuscular vitamin-A supplement treatment, ORG response demonstrated substantial recovery towards normal response characterstics, with increased response amplitudes observed at multiple retinal eccentricities.

**Figure 4:**
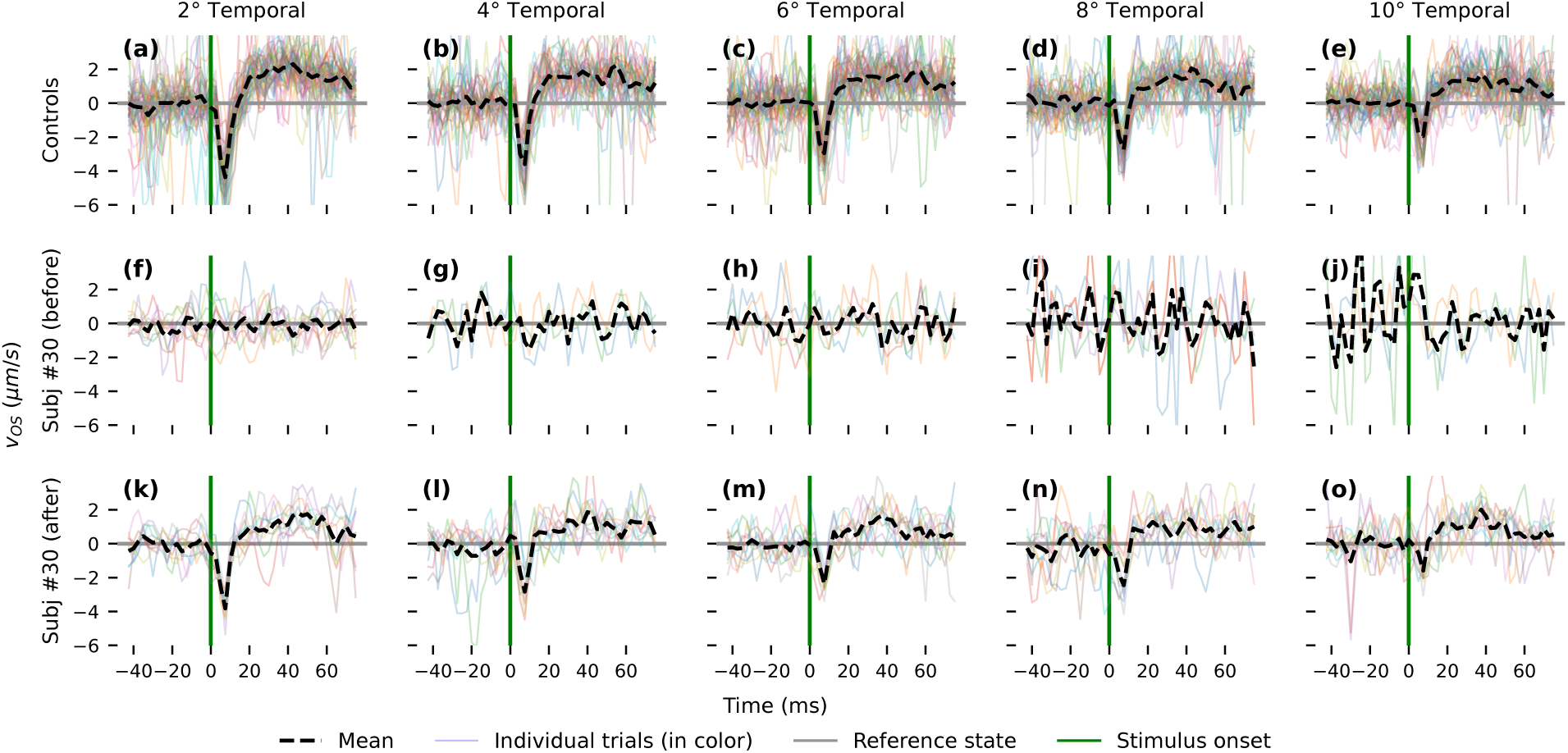
ORG responses from Vitamin-A subject vs healthy control subjects. Velocity-based ORG responses at retinal eccentricities ranging from 2*^◦^* to 10*^◦^*. (a–e) Responses from healthy controls; (f–j) responses from subject #30 with vitamin A deficiency before treatment; and (k–o) responses from the same subject after treatment. Colored traces represent individual measurements, while dashed black traces indicate the mean response at each eccentricity. The vertical green lines indicate stimulus onset, and the horizontal gray lines indicate zero velocity. Compared with healthy controls, the pretreatment responses were markedly attenuated, whereas posttreatment measurements showed recovery of the characteristic early contraction and subsequent elongation components.

Quantitative analysis of ORG parameters demonstrated reduced values of both *v_min_*and *v*_20−40_ prior to treatment compared with healthy control subjects (Fig. 5). Following treatment, both parameters increased toward values observed in healthy controls, indicating recovery of photoreceptor functional responses after vitamin-A supplementation.

**Figure 5:**
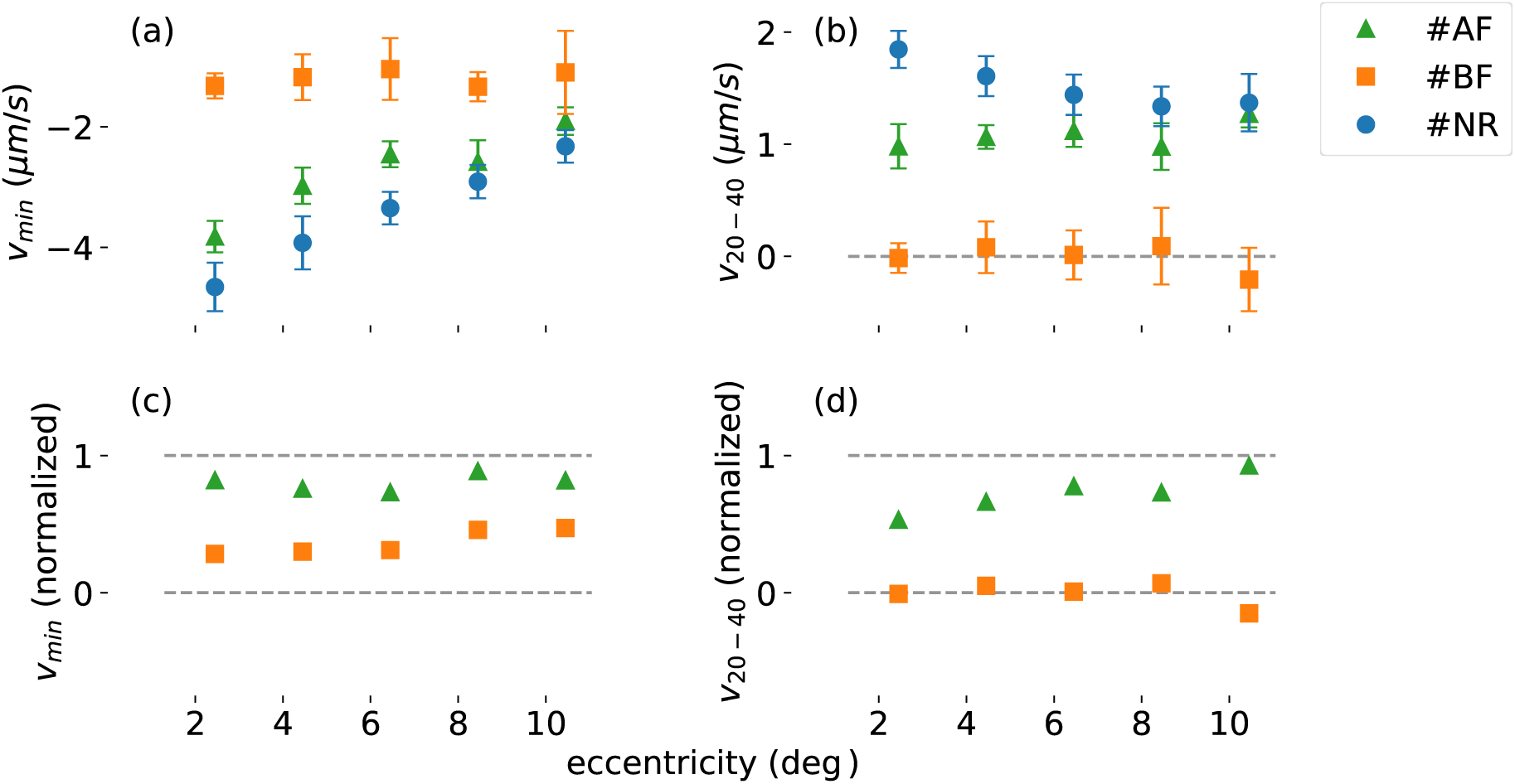
ORG parameters from a vitamin-A deficient subject before and after treatment. Comparison of ORG parameters before and after vitamin-A supplementation. (a) *v_min_*, (b) *v*_20_*_−_*_40_, (c) normalized *v_min_*, and (d) normalized *v*_20_*_−_*_40_ across retinal eccentricities. Prior to treatment, both parameters were substantially reduced compared with healthy control values. Following treatment, ORG parameters increased toward normal values across retinal eccentricities.

### 4.4 ORG responses in unexplained vision loss

Subjects with unexplained vision loss (#27 and #28) exhibited abnormal ORG parameters compared with healthy control subjects (Fig. 6). While *v_min_*values remained relatively preserved across retinal eccentricities, substantial attenuation of *v*_20−40_ was observed in both subjects. In several retinal locations, normalized *v*_20−40_ values approached or fell below zero, indicating markedly attenuated outer segment elongation responses. These findings demonstrate that ORG parameters can detect functional abnormalities in subjects with unexplained vision loss despite relatively preserved early contraction responses.

**Figure 6:**
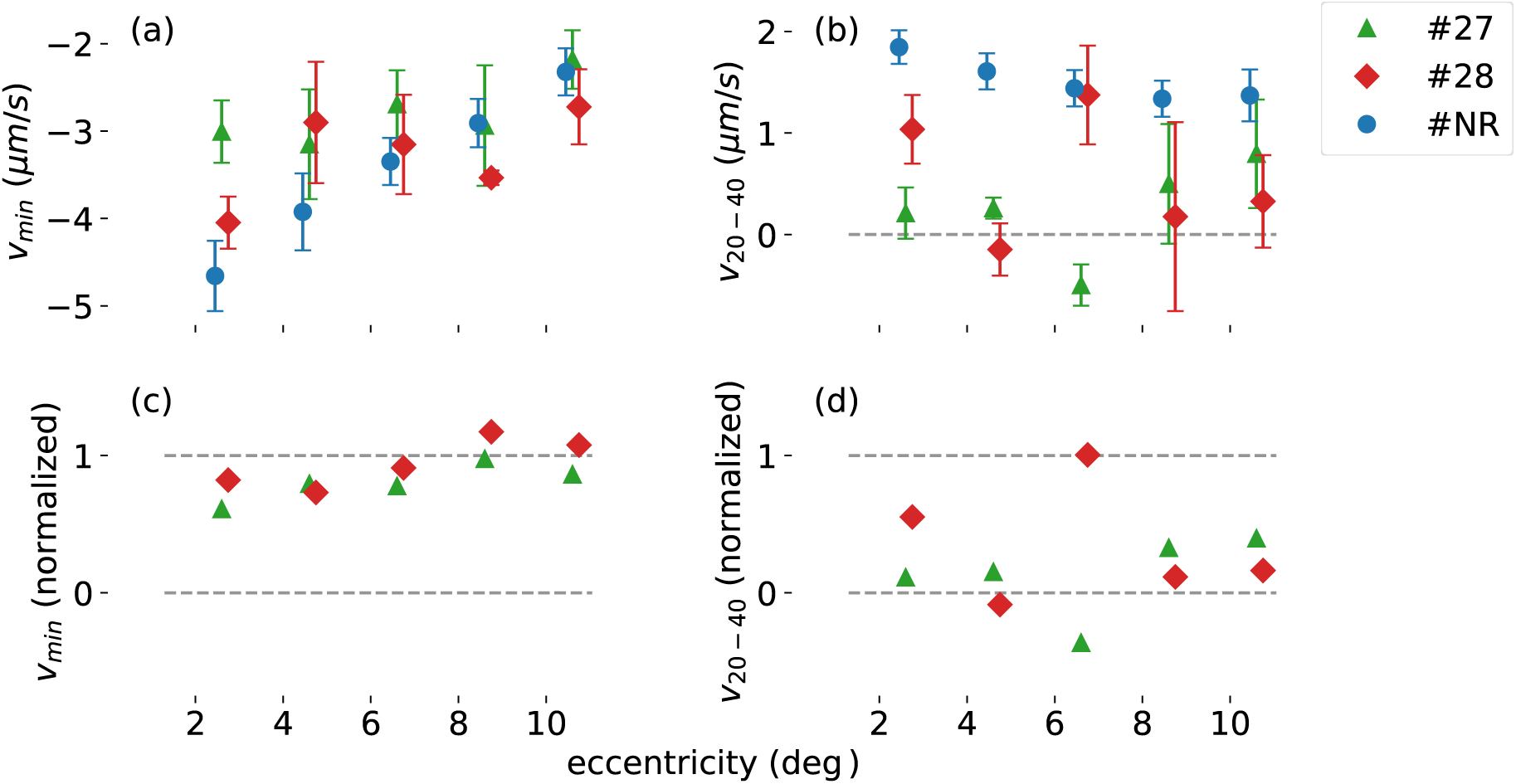
ORG parameters in subjects with unexplained vision loss. While *v_min_*values remained relatively preserved, substantial attenuation of *v*_20_*_−_*_40_ was observed across retinal eccentricities in both subjects.

### 4.5 ORG responses in diabetic subjects

Subjects with diabetes (#60 and #61) exhibited abnormal ORG parameters compared with healthy control subjects (Fig. 7). While *v_min_*was relatively preserved across the retinal eccentricities, attenuation of *v*_20−40_ was observed in both subjects. Notably, reduced *v*_20−40_ is observed in the subject without clinically visible retinopathy (#61). These observations suggests that ORG may detect photoreceptor functional abnormalities in diabetic subjects prior to the appearance of clinically detectable retinal pathology.

**Figure 7:**
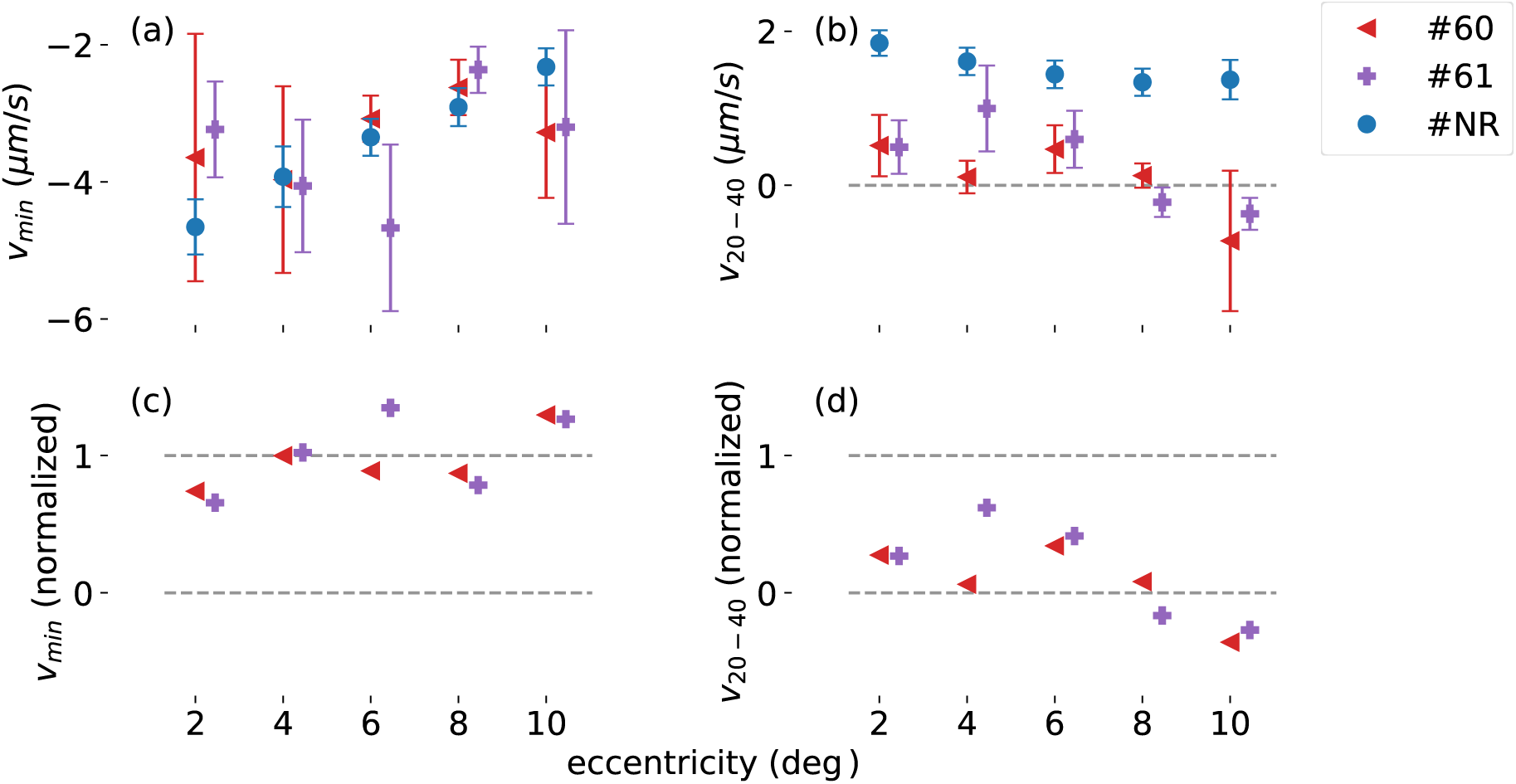
ORG parameters in diabetic subjects. Subject #60 exhibited diabetic retinopathy, while subject #61 had diabetes without clinically visible retinopathy. While normalized *v_min_* values remained relatively preserved, attenuation of normalized *v*_20−40_ values was observed in both subjects, including the subject without clinically visible retinopathy.

### 4.6 ORG responses in inherited retinal degeneration

ORG responses from two subjects with retinitis pigmentosa (subjects #33 and #34) and one subject with Stargardt maculopathy (subject #23) were compared with the healthy control cohort across retinal eccentricities from 2*^◦^* to 10*^◦^* (Fig. 8). All three inherited retinal degeneration subjects exhibited reduced ORG responses relative to healthy controls. The minimum velocity (*v*_min_) was consistantly smaller than that of the healthy cohort across most eccentricities, indicating attenuation of the early response (Fig. 8a). Likewise, the late recovery parameter (*v*_20−40_) was substantially reduced compared with controls and remained close to zero at all eccentricities (Fig. 8b), whereas healthy subjects exhibited positive values throughout the measured retinal locations. After normalization to the healthy control mean, all inherited retinal degeneration subjects demonstrated reduced responses in both parameters (Fig. 8c, d). Although inter-subject variability was present, both retinitis pigmentosa and Stargardt maculopathy showed a similar trend of diminished ORG function relative to healthy controls, consistent with impaired photoreceptor function in these inherited retinal diseases.

**Figure 8:**
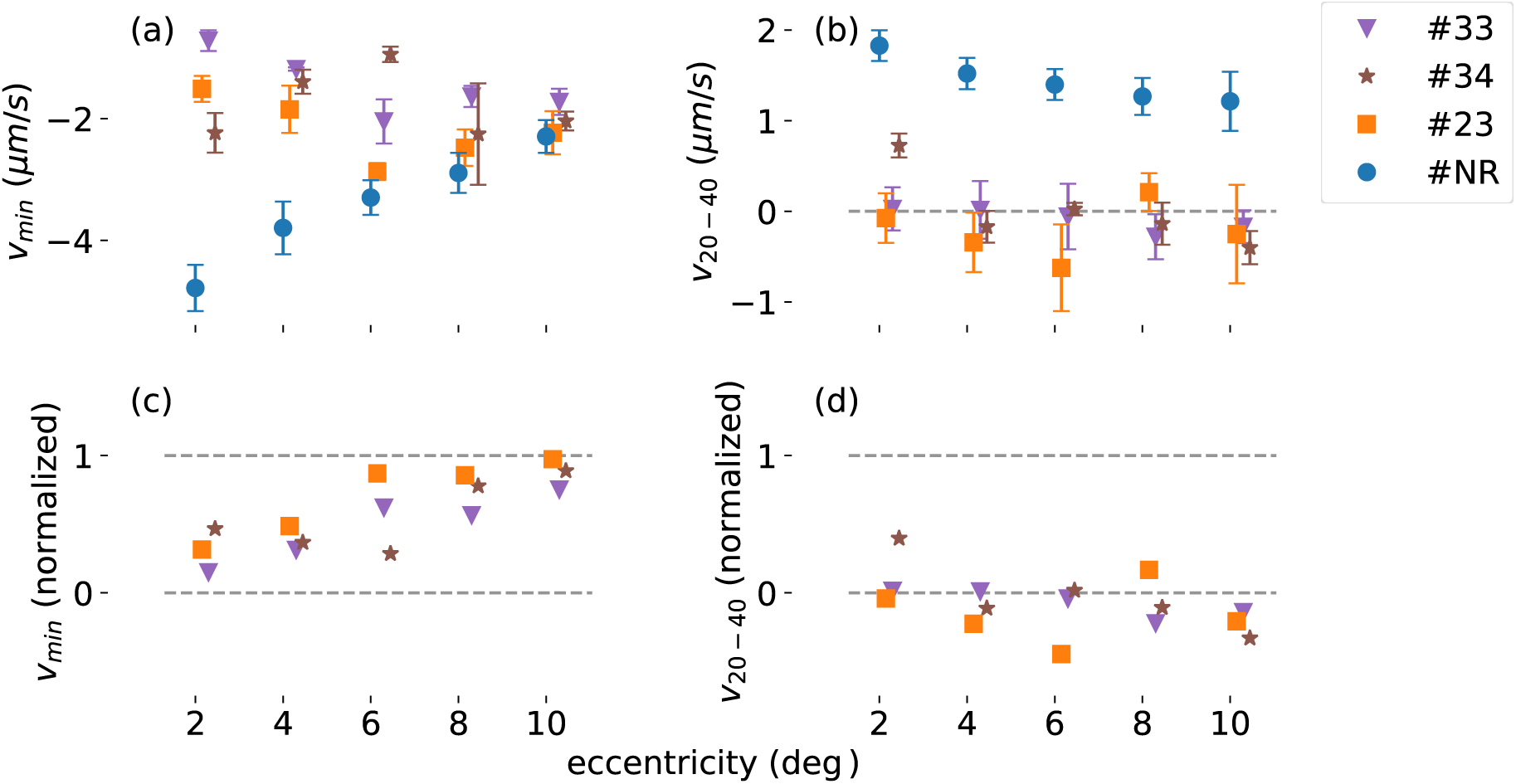
ORG parameters in inherited retinal degeneration compared with healthy controls. ORG responses from two subjects with retintis pigmentosa (subjects#33 and #34) and one subject with Stargardt maculopathy (subject#23) were compared with the healthy control cohort (#NR) at retinal eccentricities from 2*^◦^* to 10*^◦^*. (a) Minimum velocity (*v*_min_). (b) Mean velocity between 20 and 40 ms after stimulus onset (*v*_20−40_). (c,d) Corresponding normalized responses relative to the healthy control mean. Both inherited retinal degeneration phenotypes demonstrated attenuated ORG responses compared with healthy controls across retinal eccentricities.

### 4.7 ORG responses in macular telangiectasia type 2

Two subjects with macular telangiectasia type 2 (MacTel Type 2) were included in this analysis (table. 1). Subjects #52 was a 62-year-old male diagnosed with MacTel Type 2, and subject #69 was a 60-year-old female diagnosed with MacTel Type 2 and type 2 diabetes without clinical evidence of diabetic retinopathy. ORG parameters from these subjects were compared with those of the healthy control cohort across retinal eccentricities ranging from 2*^◦^* to 10*^◦^* (Fig. 9). Subject #69 exhibited relatively preserved normalized *v*_min_ values across retinal eccentricities, with responses approaching those of the healthy control cohort at several locations. In contrast, both subjects demonstrated reduced normalized *v*_20−40_ values compared with healthy controls, with subject #52 exhibiting the greatest attenuation, particularly at larger retinal eccentricities (Fig. 9). These findings suggest that the elongation stage of the ORG responses was more affected than the intial contraction in subjects with MacTel Type 2.

**Figure 9:**
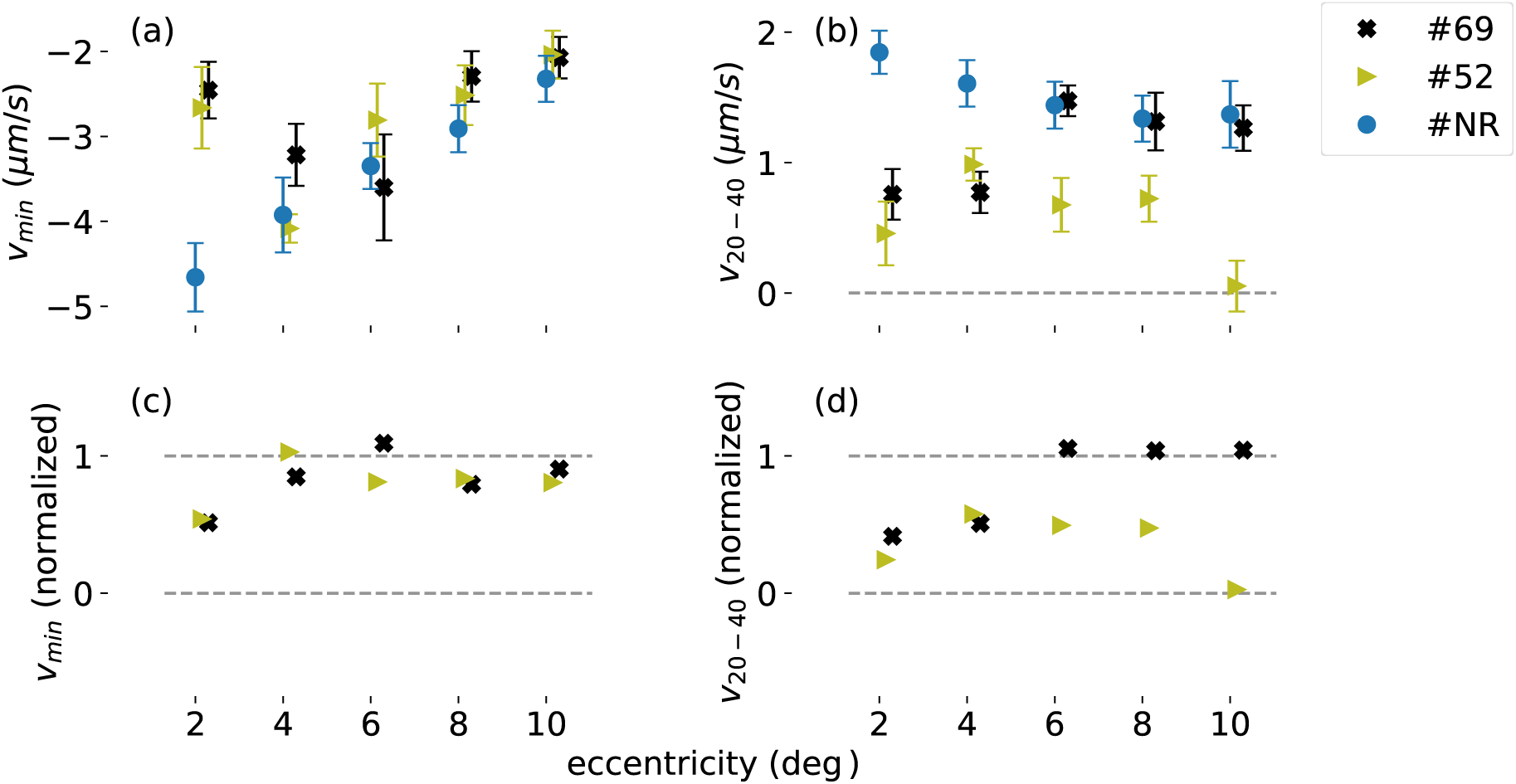
ORG parameters in subjects with macular telangiectasia Type II. ORG parameters from two subjects with macular telangiectasia type 2 (subject #52 and #69) compared with the healthy control cohort (#NR). Panels (a) and (b) show the *v*_min_ and *v*_20−40_ parameters, respectively, and panels (c) and (d) show the corresponding normalized parameters.

## 5 Discussion

### 5.1 Stages of the ORG signal and their hypothetical attributions

The optoretinographic response consists of stimulus-evoked changes in photoreceptor OS optical path length, potentially providing a direct, objective measurement of photoreceptor function. In our prior work and the prior work of other ORG investigators, the OS response seems to consist of three stages:^9^ an early contraction lasting ∼10 ms,^3, 5, 6, 25, 27^ a much larger elongation lasting hundreds of milliseconds,^3–5^ and finally a contraction toward baseline length lasting seconds.^3–5^ We have not been able to measure the last of these using the velocity-based approach, so we will focus on the first two stages. The proposed parameters *v_min_* and *v*_20,40_ are designed to quantify the rapid early contraction and slower elongation stages.

In all of the disease-affected eyes, the early contraction and/or elongation stages exhibited alterations, but the pattern of alteration varied significantly. This diversity of responses suggests that different forms of retinal dysfunction may manifest as distinct features of the ORG response. Importantly, attenuation of these responses was observed across several disease conditions, supporting the sensitivity of ORG to localized photoreceptor dysfunction. The spatially resolved nature of these measurements may therefore provide functional information complementary to conventional clinical assessments, which do not necessarily reflect photoreceptor function uniformly across the retina. The physiological basis of the ORG response remains largely unclear, although several hypotheses have been suggested.

#### 5.1.1 Early OS contraction

The early contraction of the OS has been hypothetically attributed to the early receptor potential (ERP).^6, 27^ The ERP is thought to be caused by an increase in the dipole moment along the transmembrane axis of the opsin molecule caused by photoisomerization of its chromophore.^28, 29^ In the full-field ERG, this shift in charge is visible as a small biphasic change in voltage which, interestingly, can be reversed by a second flash. The hypothetical attribution of ORG early contraction to the ERP is based on the reasoning that increased polarization of the opsin will increase the lateral repulsion of these proteins, which in turn increases the diameter of the OS discs. Due to conservation of total membrane surface area, this lateral expansion of the photoreceptor would necessarily decrease its length. Even if the ERP is not the origin of the early contraction, its time course, which has been shown by multiple investigators to initiate within 1 ms to 2 ms of stimulus onset, is consistent with that of photoisomerization and conversion to the stable intermediate meta II (or metarhodopsin II).

The clinical implication of this hypothesis, in the context of interpreting ORG responses, is that a reduction in the early contraction–i.e., a reduction in the magnitude of *v_min_* in the present work–reflects a reduction in the number of photoisomerizations in the photoreceptor. Such a reduction could be caused, for example, by low levels of unbleached photopigment in the discs. Subnormal levels of photopigment could result from dysfunction or dysregulation of the visual cycle. Of course other kinds of pathology could be implicated too. Structural changes in the photoreceptor that affect its optical properties could also reduce the amount of light absorption in the outer segment. Deleterious mutations in the opsin gene could reduce absorption as well, such as the K296E mutation, which eliminates the chromophore’s binding site^30^ or a substitution of glutamate at locus 113, which can shift the sensitivity spectrum of the opsin out of the visible range,^31^ rendering the cell insensitive to ORG stimulus sources. Other disorders could interfere with normal electrostatic effects through ultrastructural alterations in the OS discs.

#### 5.1.2 OS elongation

The time course of the OS elongation stage is consistent with it being an effect of some step(s) in the phototransduction cascade. Assuming that the initial contraction is a direct effect of photoisomerization, ORG responses lacking both contraction and elongation may be most parsimoniously attributed to lack of photoisomerization, the event that triggers the phototransduction cascade. However, when an initial contraction is present and elongation is absent, phototransduction steps downstream of photoisomerization are naturally implicated. A transducin knockout mouse model was shown to lack light-evoked OS elongation, leading to the suggestion that dissociation of a transducin subunit into the aqueous phase of the cytosol caused an osmotic shift of water into the cell.^16^ Under this view, abnormalities in ORG elongation could be attributed to mutations in transducin such as GNAT1. The same authors revised their hypothesis, and later attributed the osmotic pressure partly to phosphate dissociated from phosphodiesterase (PDE) during activation.^32^ This hypothesis is consistent with the transducin-knockout observation, since absence of transducin would also prevent PDE activation. In the same work, they show that the elongation of the OS is better explained by two mechanisms with different time constants, and claim that the second component is the movement of water out of the cytosol into the disc membranes, via the opsin protien.^32^ The hypothesis that OS elongation is due to multiple mechanisms was confirmed by a separate study.^9^ A third mechanism for OS elongation was proposed after observations of the oscillatory response of OS length to flicker stimuli.^33^ These investigators attributed OS elongation to mechanical separation of the discs by cross-links between corresponding PDE proteins on adjacent discs. The phosphate osmolarity hypothesis and the cross-linked PDE hypothesis lead to overlapping hypothetical scopes, and are not mutually exclusive.

In AO-ORG studies,^3, 4^ the highest rate of OS elongation appeared to occur between 20 ms to 40 ms post-stimulus. Moreover, in our earlier investigation of the velocity-based ORG method, we observed that positive values of *v_OS_* were most reliably found in the same interval. As such, we believe that the *v*_20−40_ ORG parameter should reflect the rate of elongation. While not discussed directly, it is evident from other studies that the rate of OS elongation is related to the maximum OS length.^5, 6^ In our study of the rod ORG, we found that the rate of elongation was more closely associated with stimulus dose than the maximum excursion of the OS.^8^

#### 5.1.3 Eccentricity dependence of contraction and elongation stages

We and others have reported an eccentricity dependence^25, 34^ of both initial contraction and elongation components, possibly mediated by eccentricity-dependence of OS length. While the present work does not analyze topographic variation in response (i.e., in relation to lesions), the eccentricity-dependence of the responses must be taken into account before inferring any disease-related attenuation of the signal.

### 5.2 Hypothetical interpretation for disease-related changes in ORG response

#### 5.2.1 Vitamin A deficiency

Vitamin A deficiency provides a particularly informative case with which to interpret alterations in the ORG response because vitamin A is required for the formation and regeneration of the chromaphore that is photoisomerized. Prolonged deficiency can reduce functional opsin, impair visual sensitivity, and eventually lead to structural changes in photoreceptor outer segments.^35^ Previous electrophysiological studies have demonstrated markedly impaired rod and cone function in vitamin A deficiency, with substantial recovery following vitamin A supplementation.^35, 36^ McBain *et al.* reported recovery of rod and generalized cone function within days of vitamin A administation, while Poornachandra *et al.* similarly observed rapid improvement within one week and near-normal ERG response by one month.^35, 36^

In the present study, subject #30 exhibited markedly attenuated ORG responses before treatment, followed by recovery accompanied by improved visual acuity from 20/150 to 20/30, providing concordant evidence of improved visual function following treatment.

These observations are particularly relevant to the proposed mechanisms of the two ORG components. If the early OS contraction is an indicator of the number of photopigment molecules undergoing photoisomerization,^27^ depletion of available visual pigment during vitamin A deficiency would be expected to attenuate this component. Restoration of the early response following supplementation would therefore be consistent with recovery of photopigment availability. The attenuation and subsequent recovery of the *v*_20−40_ response may likewise be consistent with restoration of downstream phototransduction-dependent processes, given the proposed association between OS elongation and activation of the phototransduction cascade.^16^ Experimental evidence also indicates that vitamin A deficiency alters phototransduction-related pathways and compromises visual function.^37^ However, because the molecular origins of both ORG components remain incompletely established, these interpretations should be regarded as mechanistic hypotheses rather than direct evidence of the underlying processes.

The treatment-associated recovery observed here is consistent with previous ERG studies demonstrating that retinal dysfunction associated with vitamin A deficiency can be substantially reversible following supplementation.^35, 36^ More recent clinical evidence similarly demonstrates improvement in visual symptoms and retinal findings following vitamin A repletion.^38^ Whereas ERG provides an integrated electrophysiological assessment of retinal function, the present ORG measurements demonstrate recovery of the stimulus-evoked photoreceptor response with spatial resolution across retinal eccentricities. This case therefore illustrates the potential of velocity-based ORG for objectively monitoring localized photoreceptor functional changes associated with treatment.

#### 5.2.2 Unexplained vision loss

Subjects#27 and #28, both of whom presented with unexplained vision loss, exhibited a relatively preserved *v*_min_ (≥60 % of normal everywhere, and ≥85 % in most loci). Fig. 6 shows that subject #27 exhibits clear contraction at all eccentricities. On the other hand, in both subjects, *v*_20−40_ was attenuated across most retinal eccentricities. Example ORG responses from subject #27 show a clear early contraction, whereas the subsequent elongation is markedly reduced or absent at most locations. A preliminary analysis of these subjects was presented previously.^22, 39–41^ Under the hypotheses discussed above, the relative preservation of *v*_min_ may indicate that the processes responsible for the early contractile response remain at least partially intact. If this component is associated with photoisomerization, the observed pattern would be consistent with relatively preserved photopigment. In contrast, attenuation of *v*_20−40_ may reflect impairment of processes downstream of photoisomerization, consistent with the proposed dependence of OS elongation on activation of the phototransduction cascade. The ORG abnormalities were accompanied by abnormalities in conventional clinical testing, although the cause of vision loss remain unresolved in both subjects. Subject#27 had bilateral vision loss and substantially reduced ERG responses in both eyes. In subject#28, full-field ERG suggested impairment of rod bipolar-cell and cone function, with clinical report noting a missing electronegative (B-wave) wavefrom. Genetic testing has not been performed in subject #27, whereas variants identified in subject#28 were not considered clinically relevant to the observed vision loss. Taken together, the relative preservation of the early ORG response and attenuation of the later response in these subjects suggest that different components of the photoreceptor response may be differentially affected, even when underlysing cause of vision loss remain unkown.

#### 5.2.3 Diabetes

Although DR has traditonally been diagnosed primarily by its retinal vasular abnormalities, increasing evidence indicates that diabetes also produce neural retinal dysfunction that can precede clinically apparent vascular changes.^42, 43^ Photoreceptors may be involved in this early dysfunction, with electrophysiological studies demonstrating abnormalities in photoreceptor function in subjects with diabetes even in the absence of clinically visible DR.^44^ Subjects#60 and #61 exhibited similar patterns of ORG abnormality, with relatively preserved *v*_min_ but attenuation in *v*_20−40_ across retinal eccentricities (Fig. 7). Subject #60 had clinically apparent diabetic retinopathy, whereas subject #61 had diabetes without clinically visible retinopathy. The attenuated *v*_20−40_ in subject #61 is therefore consistent with evidence that cone dysfunction can occur in diabetes before the appearance of clinically detectable vascular retinopathy. In particular, McAnany and Park reported reduced cone phototransduction sensitivity in subjects with diabetes both with and without mild DR, while the estimated maximum photoreceptor response amplitude remained relatively preserved.^44^

Under the hypothetical attributions of the ORG response, the dissociation between the relatively preserved *v*_min_ and attenuated *v*_20−40_ may provide further insight into the functional alteration in these subjects. If the early contraction is associated with photoisomerization and the ERP, the relatively preserved *v*_min_ would be consistent with preservation of the response to the L/M-cone photopigment activation. In contrast, the attenuation of *v*_20−40_ suggests greater disruption of processes contributing to the subsequent OS elongation. Because OS elongation has been proposed to arise from events downstream of photopigment activation in the phototransduction cascade, including phototransduction-dependent osmotic charges, this pattern may reflect differential effects of diabetes on the early and later componenets of the cone ORG response.

Photoreceptors have exceptionally high metabolic demands and may be particularly susceptible to metabolic disturbances associated with diabetes.^45^ Experimental studies in diabetic animals have implicated photoreceptor oxidative stress, mitochondrial dysfunction, and inflammatory signaling as possible contributors to diabetic retinal dysfunction.^42, 45^ These mechanisms could provide a biological basis for altered photoreceptor function in diabetes, although the present ORG measurements do not identify the molecular process responsible for the attenated elongation response. Taken all together, the ORG observations in these two subjects are consistent with the growing evidence that photoreceptor dysfunction can occur in diabetes and not be secondary to clinically detectable vascular abnormalities. The observation of attenuated later ORG response in subject#61, despite the absence of clinically visible retionpathy, is therefore provide complementary information about localized cone photorectptor function in diabetic retinal disease, including functional alterations that are not apparent from the vascular features used to clinically classify DR.

#### 5.2.4 Inherited retinal degenerations

Reduced sensitivity of cone (and rod) photoreceptors is observed in Stargardt’s disease,^46^ which is consistent with the attenuated ORG responses observed above (Fig. 8). The disease is caused by a mutation in the ABCA4 gene, which encodes a flippase required for removing the isomerized chromophore all-trans retinal from the outer segment disc.^47^ All-trans retinal and some of its derivatives are cytotoxic and thus it must be cleared from the outer segment. Impairment of the ABCA4 protein causes accumulation of these materials. There is evidence that photoreceptor dysfunction can precede outright atrophy of the RPE cells,^48^ which may suggest that accumulation of bisretinoids in the outer segment may interfere directly with phototransduction. It is also known that Stargardt’s disease delays dark adaptation, possibly by slowing of the visual cycle due to impairment of the RPE,^49^ due to accumulation of the bisretinoid N-retinylidene-N-retinylidene ethanolamine (A2E). A2E causes several kinds of dysfunction in RPE cells including impaired phagocytosis–a homeostatic process necessary for photoreceptor health.^50^ We have separately shown that the amplitude and kinetics of the ORG response depend on photopigment replenishment,^51^ so delayed photopigment regeneration may explain the observed attenuation of response. Under this view, the reduction in photoreceptor sensitivity may be secondary to RPE dysfunction. Rod photopigment is regenerated through a multi-step enzymatic pathway involving RPE cells. Cone photopigment is also regenerated via the RPE pathway, but cones engage in a second visual cycle via the Müller glia, but the relative contribution of the latter is thought to be smaller than that of the RPE visual cycle.^52^

In the Stargardt’s patient (#23), there appears to be an attenuation of the early contraction in the parafovea (2*^◦^* to 4*^◦^*). The elongation stage is largely absent, even–interestingly– when the early contraction appears normal. This suggests that Stargardt’s disease impacts the phototransduction process independently of its effects on the mechanisms of early contraction. Even if the disease negatively impacts photopigment levels by hampering the visual cycle, direct effects in the photoreceptor seem likely.

Disease-related changes in *v*_20−40_ are by themselves non-specific, as upstream abnormalities (e.g., in photopigment availability) would affect it. Patient #23, who has Stargardt maculopathy with a pathogenic mutation in ABCA4, has *v*_20−40_ values significantly lower than the control subject’s at all eccentricities (see Fig. 8(d)). However it would be incorrect to assume that this indicates a phototransduction failure because we also observed attenuated *v_min_* at most locations, as low as 33% of normal at 2 deg and 54% of normal at 4 deg (Fig.8(c)). Assuming that photoisomerization drives *v_min_*, we can infer that the patient’s opsins are absorbing abnormally low numbers of photons. This could be due to any number of factors–visual cycle defects, outer segment structural flaws, or even prereceptoral factors such as media opacity or cataract (although those were ruled out clinically). It is interesting to note that the abnormal ORG findings are consistent with the patient’s subjective tests, but that neither ERG nor mfERG detected any abnormalities. The spatial resolution of the mfERG (∼1 deg) is similar to ours (1.2 deg), which is of special interest. This could be due to noise or error in one or both modalities, but if not it suggests adaptation of the cones to subnormal pigment isomerization and early phototransduction, possibly through modulation of phototransduction gain.

RP comprises a genetically heterogeneous group of inherited retinal degenerations in which rod photoreceptor degeneration is typically followed by secondary cone dysfunction and loss. Cone dysfunction may arise from alterations in the retinal environment associated with RPE degeneration and impaired support of photoreceptor homeostasis. An additional proposed mechanism involves reduced availability of rod-derived cone viability factor (RdCVF), which may compromise cone metabolism and survival.^53, 54^

Previous ORG investigations have demonstrated impaired cone responses in RP, including retinal regions with relatively preserved cone density.^12^ Wendel et al. further reported reduced ORG responses in RP despite relatively preserved visual sensitivity in some subjects.^13^ More recently, Liu et al. reported longitudinal changes in ORG responses that exceeded changes in conventional OCT-derived structural measurements in patients with RP.^55^ Together, these studies highlight the potential of ORG to provide functional information complementary to conventional retinal assessments.

In the present study, both subjects with RP(#33 and #34) exhibited attenuated lightevoked photoreceptor OS responses compared with healthy controls (Fig. 8). Although the initial contraction component (*v*_min_) remained measurable, the later elongation component (*v*_20−40_) was substantially attenuated in both subjects.

While the genetic cause of RP in these patients is unknown, most forms of RP affect rods before cones. These alterations in cone responses may reflect secondary cone dysfunction following rod degeneration, potentially associated with RPE-mediated changes in photoreceptor support or reduced RdCVF availability. However, without information about the underlying genetic mutations, the specific mechanisms responsible for the observed ORG abnormalities cannot be determined. The ability to characterize these responses using clinical verlocity-based ORG extends previous observations obtained primarily with adaptive optics-based ORG approaches.

#### 5.2.5 Macular telangectasia type 2

MacTel Type 2 is a progressive retinal disorder characterized by neurodegenerative and vascular abnormalities, typically beginning in the temporal parafoveal region.^56, 57^ Histopathological studies have demonstrated depletion of Müller glial cells and abnormalities of the photoreceptor layers in affected retinal regions.^58^ Adaptive optics imaging has further revealed disruption of the cone photoreceptor mosaic and reduced parafoveal cone density, associated with decreased retinal sensitivity and abnormalities of the photoreceptor IS/OS junction on OCT.^59^ Alterations in serine and sphingolipid metabolism, including the accumulation of potentially neurotoxic deoxyspingolipids, may contribute to photoreceptor dysfuction.^60, 61^

In the present study, the two subjects with MacTel Type 2 exhibited alterations in the light-evoked photoreceptor OS response, particularly in the later elongation component (*v*_20−40_). The intial contraction component (*v*_min_) was relatively preserved in some measurements, and the two components of the ORG response appear to be affected diferently. Although the limited number of subjects preculudes conclusions regarding disease-specific response patterns, these preliminary observations demonstrate the feasibility of detecting and characterizing altered photoreceptor responses in eyes with MacTel Type 2 using clinical velocity-based ORG.

The observed alterations may be considered in the context of the underlying photoreceptor and Müller glial abnormalities associated with MacTel Type 2. Previous psychophysical investigations have demonstrated impaired cone pigment regeneration and elevated visual thresholds in MacTel Type 2,^62^ providing independent evidence of cone functional abnormalities. Although Müller glial dysfunction has been proposed to contribute to impaired cone visual pigment regeneration, the relationship between these abnormalities and the millisecond-scale velocity-based ORG response remain uncertain. Delayed pigment regeneration observed over minutes cannot directly explain the altered ORG velocities measured within tens of milliseconds. However, it remains possible that the standard 5 min dark adaptation period in the experimental protocol leaves MacTel Type 2 patients with a reduced complement of unbleached photopigment, as compared to normal subjects.

Furthermore, subject#69 also had T2D without clinically apparanet diabetic retinopathy. Previous electrophysiological studies have identified cone functional abnormalities in diabetes even in the absence of visible retinopathy,^44^ suggesting that diabetes may represent additional factor influencing photororeceptor function. Therefore, the relative contibution of MacTel Type 2 and diabetes to the observed ORG abnormalities cannot be distinguished in this subject. Collectively, these preliminary observations demonstrate the capablity of clinical velocity-based ORG to detect and characterize altered light-evoked photoreceptor responses in subjects with MacTel Type2. This approach offers the potential to provide complementary function information alongside conventional structural OCT imaging.

### 5.3 Potential limitations

The present study demonstrates the capability of clinical velocity-based ORG to characterize light-eveoked photoreceptor responses across a range of retinal diseases. Nevertheless, several limitations should be considered.

First, the number of subjects within each disease group was small, limiting our ability to establish disease-specific response patterns. Although patient responses were compared with those of six healthy subjects, the control cohort was relatively small and generally younger than the patient cohort. Consequently, the potential contribution of age-related variability to observed differences cannot be excluded. Larger, age-matched cohorts would help establish normative ranges and characterize the variability of velocity-based ORG parameters.

Second, the present velocity-based ORG approach requires identifiable photoreceptor IS/OS and COST reflection to measure changes in outer-segment optical path length. In eyes with advanced photoreceptor degeneration, these layers may be severely disrupted or absent, preventing reliable segmentation and ORG signal extraction. Subjects in whom the required photoreceptor layers could not be identified were therefore excluded from the reported analysis. Consequently, the present results primarily reflect eyes with sufficient residual photoreceptor structure to support ORG measurements.

Third, measurement reliability is sensitive to involuntary eye and body movements and fixation instability. In some subjects excluded from the current study, particularly older individuals exhibiting tremor or shivering, substantial motion-related noise prevented reliable extraction of ORG responses. In other excluded subjects, poor fixation limited the ability to obtain stable measurements at the intended retinal locations. Residual eye motion, phase noise, and segmentation variability can further affect the estimated velocities.^23^ Improvements in motion correction, acquisition stability, and signal-quality assessment may increase the applicability of the method to patients with impaired fixation or involuntary movements. Fourth, the physiological origions of the early contraction and later elongation components remain incompletely understood. Alterations in *v*_min_ and *v*_20−40_ therefore cannot be uniquely attributed to specific molecular or cellular mechanisms, phototransduction, or outer segment integrity may contribute to the observed responses. These parameters should be interpreted as complementary descriptors of photoreceptor function rather than direct measures of individual physiological processes.

Finally, although longitudinal ORG measurements were obtained in the subject with vitamin A deficiency, the present study was primarily designed to demonstrate the capability of clinical velocity-based ORG to detect and characterize photoreceptor functional abnormalities across different retinal diseases, rather than to establish diagnostic accuracy or systematically assess disease progression.

## 6 Conclusion

This study demonstrates that clinical velocity-based optoretinography (ORG) can quantitatively characterize the early contraction and later elongation components of light-evoked photoreceptor responses, revealing functional abnormalities across a spectrum of retinal diseases. By enabling objective, noninvasive functional assessment with clinically compatible instrumentation, this approach extends ORG beyond specialized systems and provides functional information complementary to conventional structural retinal imaging. These findings establish a foundation for the clinical translation of ORG as a potential quantitative biomarker for assessing photoreceptor dysfunction. Future studies involving larger, age-matched and multi-center cohorts, comprehensive normative databases, and longitudinal assessments will help establish its diagnostic and prognostic value and support its integration into routine clinical ophthalmic imaging.

## Funding

NIH grants R01-EY-034340, R01-EY-033532, R01-EY-031098, R01-EY-26556, and P30-EY-012576.

## Acknowledgments

We would like to thank: Susan Garcia and Michael Lien (UC Davis) for their efforts in patient coordination and Profs. John S. Werner, Paul A. Sieving, and Ala Moshiri (UC Davis) for helpful discussions about photoreceptor function and disease; Sricharan Pusarla for assisting in data management; and Prof. Yifan Jian (OHSU) for development of OCT acquisition software.

## Disclosures

The authors declare no conflicts of interest.

## Data Availability Statement

The datasets generated and analyzed during the current study are not publicly available at this time but are available from the authors upon request.

## Notes

### Competing Interest Statement

The authors have declared no competing interest.

